# Core binding factor leukemia hijacks T-cell prone PU.1 antisense promoter

**DOI:** 10.1101/2020.05.29.120857

**Authors:** E. van der Kouwe, G. Heller, A. Czibere, L.H. Castilla, R. Delwel, A. Di Ruscio, A.K. Ebralidze, M. Forte, L. Kazianka, C. Kornauth, T. Le, K. Lind, I.A. Monteiro Barbosa, A. Pichler, J.A. Pulikkan, A-S Schmolke, H. Sill, W.R. Sperr, A. Spittler, B. Q. Trinh, P. Valent, K. Vanura, R.S. Welner, J. Zuber, D.G. Tenen, P.B. Staber

## Abstract

The blood system serves as a key model for cell differentiation and cancer. It is orchestrated by precise spatiotemporal expression of the hematopoietic master regulator PU.1^1–4^. PU.1 gene expression is regulated through enhancer-promoter interactions within a topologically associated domain (TAD)^5,6^. PU.1 levels increase during myeloid differentiation while failure to do so results in myeloid leukemia^7^. In contrast, T-cell differentiation requires PU.1 to be completely switched off^8–10^. Little is known about the precise mechanisms of PU.1 repression, physiological as in T-cell differentiation, or pathological as in leukemia. Here we demonstrate that the down-regulation of PU.1 mRNA is a dynamic process involving an alternative promoter^11^ in intron 3 that is induced by RUNX transcription factors driving noncoding antisense transcription. Core binding factor (CBF) fusions, RUNX1-ETO and CBFβ-MYH11 in t(8;21) and inv(16) acute myeloid leukemia (AML)^12^, activate the PU.1 antisense promoter, thus shifting from sense towards antisense transcription and blocking myeloid differentiation. In patients with CBF-AML, we found that an elevated antisense/sense ratio represents a hallmark compared to normal karyotype AML or healthy CD34+ cells. Competitive interaction of the enhancer with the proximal or the antisense promoter are at the heart of differential PU.1 expression during myeloid and T-cell development. Leukemic CBF fusions thus utilize a physiologic mechanism employed by T-cells to decrease sense PU.1 transcription. Our results identify the first example of a sense/antisense promoter competition as a crucial functional switch for gene expression perturbation by oncogenes. This novel basic disease mechanism reveals a previously unknown Achilles heel for future precise therapeutic targeting of oncogene-induced chromatin remodeling.

## MAIN

To examine the developmental activation of the PU.1 (*SPI1*) locus (**Fig.1a**) throughout all human hematopoietic differentiation stages (**Fig.1b**), we analyzed publicly available data identifying open chromatin regions using the assay for transposase-accessible chromatin with high-throughput sequencing (ATAC-seq)^13^. We identified highly versatile cell type specific accessibilities at a conserved region in intron 3 that we previously identified as a promoter of a long noncoding antisense transcript (asRNA)^11^ (**Fig.1c-d**). Chromatin accessibility at the PrPr stayed constant compared to the antisense promoter (AsPr) during early myelopoiesis (**Fig.1c**) whereas in mature CD4+/CD8+ T-lymphoid cells these sites were absent (**Fig.1d**). For in-depth analysis of sense (mRNA) and antisense (asRNA) transcript levels, we performed compartment enriched Northern Blot and strand-specific reverse-transcription quantitative polymerase chain reaction (RT-qPCR) quantification assays (**Extended Data Fig.1a-f**) and RT-qPCR on sorted bone marrow progenitor and mature peripheral blood cells (**Extended Data Fig.1g-h, Extended Table1**). Quantification of chromatin accessibility and transcript expression revealed that during myelopoiesis, progenitors and differentiated cells constantly displayed higher PrPr accessibility and mRNA transcription compared to AsPr accessibility and asRNA expression (**Fig.1e**). In contrast, during lymphopoiesis, we found high AsPr accessibility and asRNA expression in lymphoid-primed multipotent progenitor (LMPP) and common lymphoid progenitor (CLP) populations which preceded locus shutdown in T cells (**Fig.1f**). Ratio calculations of AsPr/PrPr accessibility and asRNA/mRNA expression revealed a constant ratio for PU.1 promoters and transcripts during myelopoiesis (**Fig.1g**) but a striking AsPr and asRNA increase during lymphopoiesis reaching a peak before complete repression in T-cells (**Fig.1h**). Promoter accessibility ratios and transcript expression ratios demonstrated a strong correlation for each population (**Fig.1i**). Increased AsPr/PrPr accessibility ratios (**Fig.1j**) and increased asRNA/mRNA transcript ratios (**Fig.1k**) distinguished lymphoid from myeloid populations during blood cell differentiation. Thus, the ratio of antisense/sense transcription indicates cellular fate in hematopoiesis. To investigate the mechanisms that drive asRNA transcription, we investigated expression levels of hematopoietic transcription factors during thymic differentiation (**Fig.1l**). Using transcript sequencing (RNA-seq) of thymic progenitor cells^14^, we identified that increasing RUNX factor expression matched with decreasing PU.1 expression in early thymic progenitor (ETP), CD1a-pro-T and CD1a+ pro-T cells (**Fig.1m-o, Extended Fig.1i-j**). In addition, a consensus RUNX binding motif was located in PU.1 AsPr (**Fig.1p**) which specifically bound RUNX1 as well as RUNX3 in electromobility shift assays (EMSA) using extracts from Jurkat and HL-60 cells (**Fig.1q**). We next designed a luciferase reporter assay in HEK293T cells transiently transfected with reporter plasmids containing PU.1 AsPr and analyzed transactivation activity with increasing concentrations of each RUNX factor. RUNX1 and RUNX3 indeed showed dose-dependent AsPr transactivation (**Fig.1r-s, Extended Fig.1k**). RUNX factors (core binding factors) are frequently altered in human leukemia, most commonly as t(8;21)(q22;q22) or inv(16)(p13;q22) chromosomal translocations of so called core-binding factor (CBF) leukemias accounting for 15% of all AML cases^12,15^. We therefore tested the oncogenic fusion proteins RUNX1-ETO (t8;21) and CBFB-MYH11 (inv16) and found, similarly to wild type RUNX1 and RUNX3, that RUNX1-ETO and CBFβ-MYH11 fusions dose-dependently induced transactivation of PU.1 AsPr (**Fig. 1t-u**). These experiments demonstrate that PU.1 AsPr can be activated by RUNX factors and by leukemic CBF-fusion proteins.

**Fig. 1.**
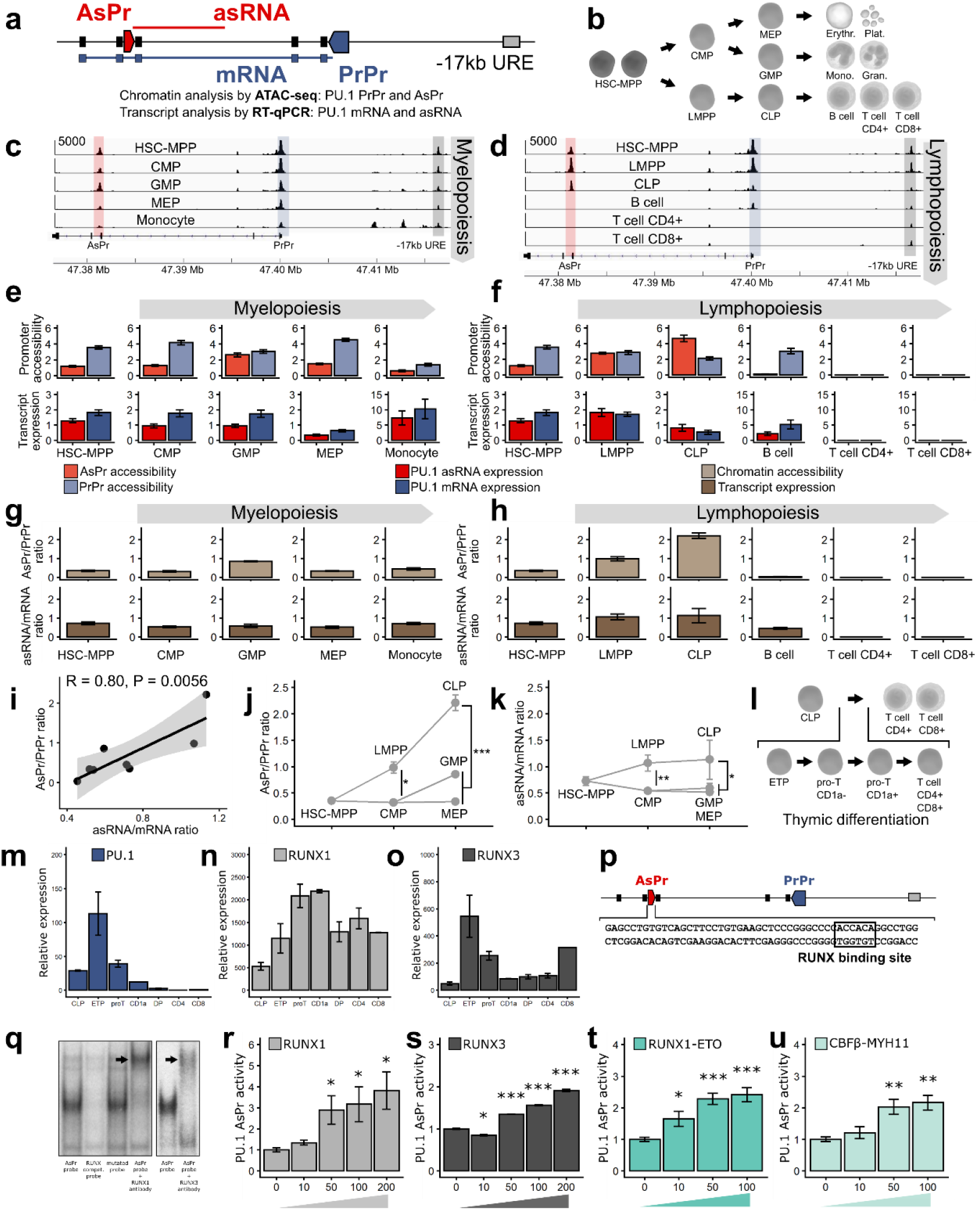
PU.1 antisense promoter activated by RUNX factors during T-lymphoid differentiation. **a**, Schematic of PU.1 locus: Proximal promoter (PrPr, blue arrow box), antisense promoter (AsPr, red arrow box), coding mRNA (blue line), antisense RNA (asRNA, red line), −17kb enhancer (upstream regulatory element, URE, grey box). **b**, Human hematopoietic cell differentiation hierarchy (HSC-MPP, hematopoietic stem cell and multipotent progenitor; CMP, common myeloid progenitor; MEP, megakaryocyte-erythroid progenitor; GMP, granulocyte-macrophage progenitor; LMPP, lymphoid-primed multipotent progenitor; CLP, common lymphoid progenitor; Erythr., erythrocyte; Plat., platelet, Mono., monocyte; Gran., granulocyte). **c-d**, Aligned reads from ATAC-seq of PU.1 locus during **c**. myelopoiesis and **d**. lymphopoiesis (RM Corces et al.). **e-f**, URE-adjusted peak quantification values for AsPr and PrPr by ATAC-seq (upper panel) and PU. 1 mRNA and asRNA (respectively dark red and dark blue) transcript profile using RT-qPCR (lower panel) for each population (mean value ± s.e.m.) in **e**, myelopoiesis (Population, replicates for ATAC-seq/RT-qPCR; HSC-MPP, n=13/9; CMP, n=8/8; GMP, n=7/9; MEP, n=7/10; Monocyte, n=6/6) and **f**, lymphopoiesis (Population, replicates for ATAC-seq/RT-qPCR; LMPP, n=3/4; CLP, n=5/4, T cell CD4+, n=5/6; T cell CD8+, n=5/6). **g-h**, Ratio of AsPr to PrPr (light brown) and asRNA to mRNA (dark brown) for each population (mean value ± s.e.m.) in **g**. myelopoiesis and **h**. lymphopoiesis. **i**, Correlation analysis of promoter chromatin accessibility ratios to transcript expression ratios. Regression line with Pearson correlation coefficient. **j-k**, Promoter and transcript ratio analysis in bone marrow hematopoietic progenitors for **j**. AsPr/PrPr chromatin accessibility ratio and **k**. asRNA/mRNA transcript expression ratio (mean value ± s.e.m.). Mann-Whitney-U test; LMPP versus CMP; CLP versus combined GMP/CMP (*P < 0.05, ***P < 0.001). **l**, Schematic of the human thymic T-lymphoid differentiation hierarchy (CLP, common lymphoid progenitor; ETP, early thymic progenitor). **m-o**, Gene expression by transcript sequencing (RNA-seq, D Casero et al.) in thymic progenitors and differentiated T cells (pro-T, CD1a-pro-T cell; CD1a, CD1a+ pro-T cell; DP, double-positive CD4+/CD8+ T cell; CD4, CD4+ T cell; CD8, CD8+ T cell) for **m**. PU.1, **n**. RUNX1 and **o**. RUNX3 hematopoietic transcription factors (mean value relative to GAPDH housekeeping gene ± s.e.m., n = 2 per population). **p**, Schematic representation of the PU. 1 locus with the antisense promoter containing a consensus RUNX binding site (black square). **q**, Gel electromobility shift assay (EMSA) with labelled PU.1 AsPr probe oligonucleotide in the presence of RUNX1 (in Jurkat cell line) or RUNX3 antibody (in HL-60 cell line) (RUNX compet. Probe, competition probe with mutated RUNX binding site). **r-u**, Luciferase reporter assays in HEK293T cells transiently transfected with PU.1 AsPr reporter plasmid and increasing **r**. RUNX1, **s**. RUNX3, **t**. RUNX1-ETO and **u**. CBFβ-MYH11 expression plasmids (plasmid concentration (ng) mean ± s.e.m., n = 4). Student T-test; 0 ng control group versus individual expression plasmid groups (*P < 0.05; **P < 0.01; ***P <0.001). We acknowledge somersault1834 (www.somersault1824.com) for providing the hematopoietic cell illustrations.

To investigate if in CBF leukemia the ratio of antisense/sense transcription is altered, we analyzed PU.1 mRNA and asRNA expression in a cohort of patient samples of normal karyotype AML (NK-AML), CBF-AML, and CD34-enriched healthy bone marrow samples. Strikingly, we found elevated PU.1 asRNA and decreased PU.1 mRNA expression resulting in a highly significant increased asRNA/mRNA ratio in CBF-AML patients (**Fig2a-c, Extended Fig.2a**). Similarly, we examined open chromatin regions of PU.1 determined in patient samples by DNaseI-sequencing (DNase-seq)^16^. Compared to NK-AML we found increased AsPr DNase hypersensitivy sites (DHS) in t(8;21) and inv(16) CBF-AML subgroups and increase AsPr/PrPr promoter DHS ratios in t(8;21) alone (**Fig2d-f, Extended Fig.2b**).To probe its functional relevance in CBF-AML we depleted PU.1 asRNA using lentiviral small-hairpin RNA (shRNA) in the t(8;21) leukemic cell line, Kasumi-1 (**Fig.2g, Extended Fig.2c-e**). PU.1 asRNA reduction lead to increased PU.1 mRNA levels (**Fig.2h**), and loss of the CD34 surface marker characteristic for immature cells (**Fig.2i**). As expected for PU.1 increase^2,3^, the cell cycle was arrested (**Fig.2j**) and cells increasingly entered apoptosis (**Fig.2k**). Importantly, PU.1 asRNA depletion was associated with a loss of in vivo leukemogenicity in a (NOD/SCID) xenograft model (**Fig.2g, l**). Cytospins of isolated targeted Kasumi-1 cells exhibited a myeloid mature differentiation phenotype compared to controls (**Fig.2m**). Other pathophysiologic alterations such as splenomegaly and bone whiting related to massive leukemic infiltration were inhibited in asRNA depleted compared to controls (**Fig.2n-o**). These data demonstrate that PU.1 antisense transcription is functionally required to establish a CBF leukemia model.

**Fig. 2.**
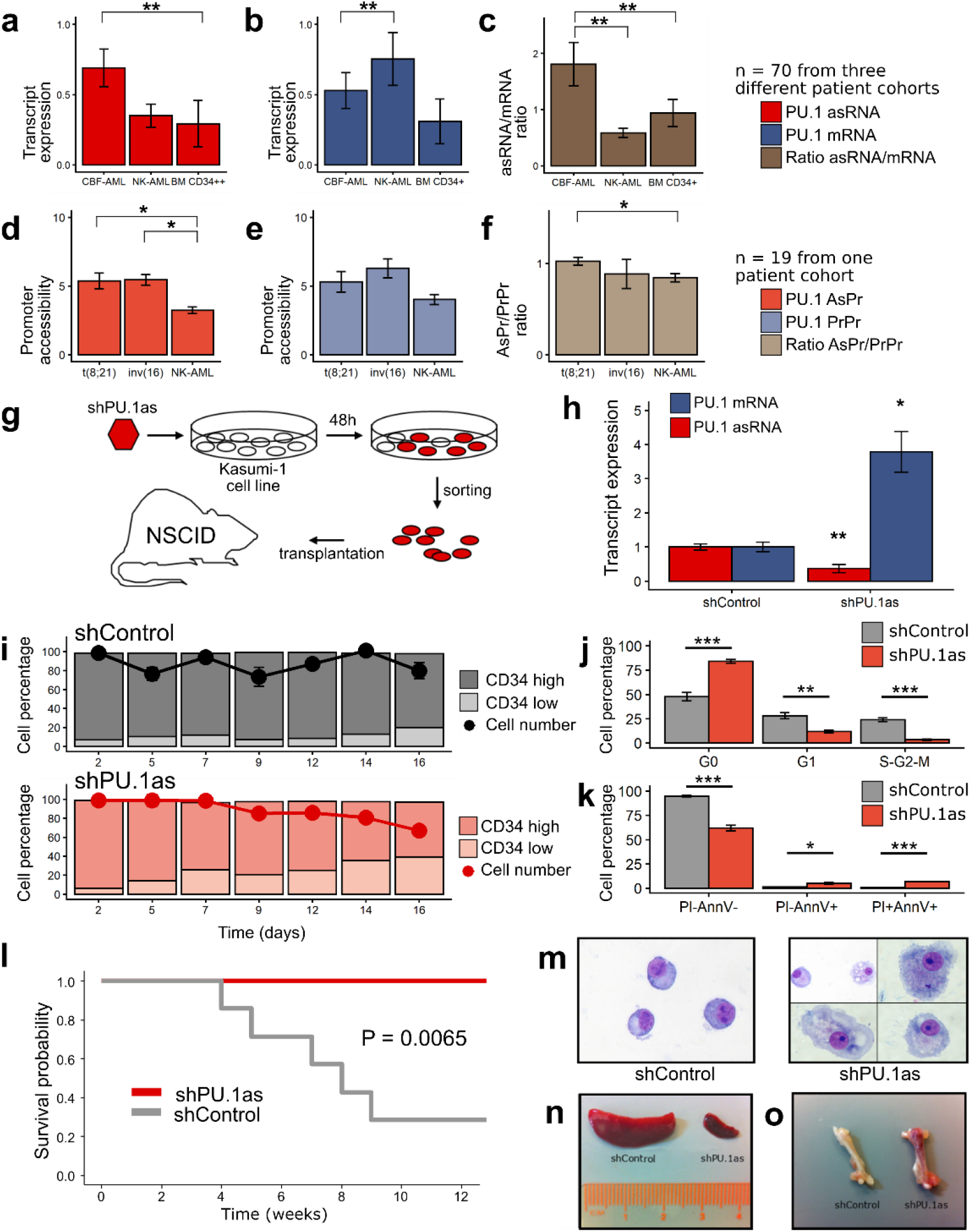
PU.1 antisense activation in core binding factor leukemia. **a-c**, Transcript quantification for **a**. PU.1 antisense (asRNA, red), **b**. mRNA (blue) and **c**. asRNA/mRNA ratio calculation in CBF-AML (n = 36), normal karyotype AML (NK-AML, n = 25) patient samples and normal CD34-enriched bone marrow (BM CD34+, n = 9) from three patient cohorts (mean value ± s.e.m.) (*P < 0.05; **P < 0.01; ns, non-significant). **d-f**, Promoter quantification for **d**. PU.1 antisense promoter (AsPr, light red), **e**. proximal promoter (PrPr, light blue) and **f.** AsPr/PrPr ratio calculation in t(8;21)-AML (n = 3), inv(16)-AML (n = 2) and normal karyotype AML (NK-AML, n = 19) patient samples (mean value ± s.e.m.) (*P < 0.05) (SA Assi et al.). **g**, Experimental workflow of shRNA knockdown of PU.1 antisense RNA (shPU.1as) in Kasumi-1 followed by fluorescence-activated cell sorting (FACS) and xenografting into immunodeficient (NSCID) mice. **h**, PU.1 mRNA and asRNA expression in Kasumi-1 (mean value ± s.e.m., n = 4) (*P < 0.05, **P < 0.01). **i**, CD34 surface marker and viability kinetics assessed by flow cytometry after shPU.1as in Kasumi-1 (mean CD34 percentage relative to day2, mean viability relative to day2 ± s.e.m., n = 4) (*P < 0.05). **j-k**, Assays for **j**. cell cycle stages after shPU. 1as and **k**. cell viability (Propodium Iodide, PI; Annexin V, AnnV; mean value ± s.e.m., n = 4) in Kasumi-1 cells (mean value ± s.e.m., shControl n = 5, shPU.1as n = 6) (*P < 0.05, **P < 0.01, ***P < 0.001). **l**, Survival probability of NOD-SCID mice xenografted with Kasumi-1 cells after shPU.1as versus shControl, (n = 7). **m**, May Grünwald/Giemsa cytospins for morphology analysis of Kasumi-1 cells. **n-o**, Leukemic cells infiltration in **n**. spleen and **o**. bone samples from mice transplanted with Kasumi-1 cells.

To further dissect the mechanism of how CBFs could drive PU.1 antisense transcription, we first depleted RUNX1-ETO using lentiviral small-hairpin RNA knockdown (shRUNX1-ETO) in t(8;21) Kasumi-1 cells (**Fig.3a-b, Extended Fig.3a-b**). Similar to PU.1 antisense depletion (**Fig.2g-h**), RUNX1-ETO depletion resulted in a loss of CD34 surface marker expression and a drop in cell viability (**Fig.3c, Extended Fig.3c-d**). Moreover, Giemsa staining of isolated targeted Kasumi-1 cells identified a mature myeloid differentiation phenotype compared to controls (**Fig.3d**). RNA-seq confirmed a switch towards myeloid differentiation after RUNX1-ETO depletion (**Fig.3e, Extended Fig.3e**) accompanied by increase of *PU.1* expression and a loss of *CD34* expression (**Fig.3f**). We next investigated PU.1 AsPr and PrPr chromatin accessibility and corresponding transcript levels in Kasumi-1 (t8;21) and ME-1 (inv16) cells after RUNX1-ETO knockdown or after specific CBFβ-MYH1 inhibition using a small molecule inhibitor of oncoprotein multimerization (AI-10-49)^17,18^, respectively. After CBF targeting, we identified decreased AsPr and relatively increased PrPr accessibility resulting in a shift towards a myeloid differentiation prone decreased AsPr/PrPr ratio in both models (**Fig.3g-h**). Similarly, CBF depletion resulted in decreased PU.1 asRNA, and increased mRNA levels (**Fig.3i**), with a decreased asRNA/mRNA ratio associated with myeloid differentiation (**Fig.3j, Fig.1g**). To directly detect strand-specific effects of CBFs on transcription we used precision nuclear run-on sequencing (PRO-seq) mapping of active RNA polymerases. Importantly, quantified PRO-seq reads at AsPr and AsPr/PrPr ratios decreased after RUNX1-ETO depletion in Kasumi-1 cells (**Fig.3k**). Noteworthy, aligned PRO-seq reads started at the previously identified RUNX binding site at the PU.1 AsPr (**Fig.3l**). These experiments demonstrate that core-binding factors cause a shift from PU.1 sense to antisense transcription.

**Fig. 3.**
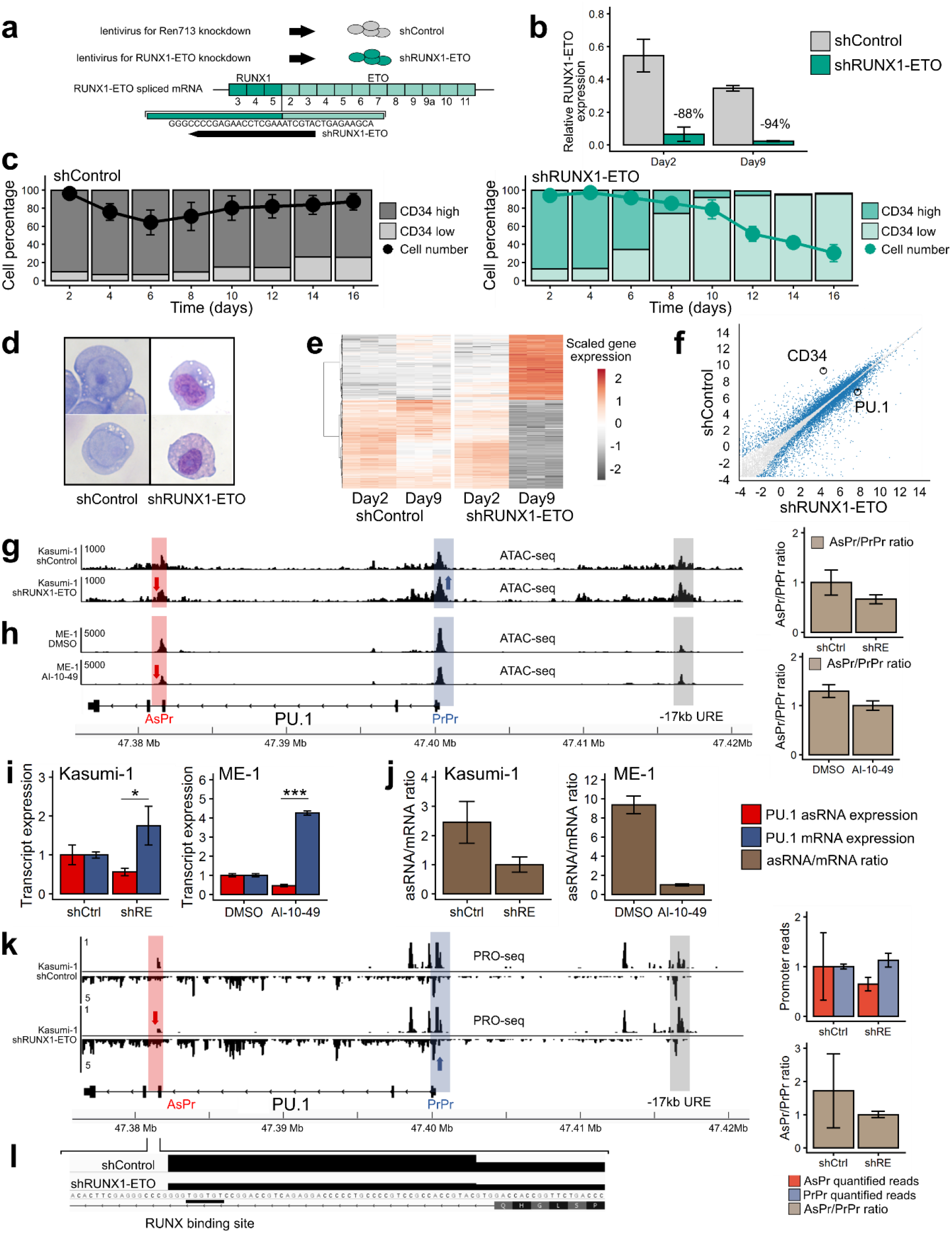
Oncogenic core binding factors shift PU.1 sense to antisense transcription. **a**, Scheme of shRUNX1-ETO knockdown design. **b**, RUNX1-ETO RT-qPCR after RUNX1-ETO knockdown at day2 and day9 in Kasumi-1 cells relative to GAPDH housekeeping gene (mean value ± s.e.m., n = 3). **c**, CD34 surface marker and viability kinetics assessed by flow cytometry after shRUNX1-ETO in Kasumi-1 cells (± s.e.m., n = 2). **d**, May Grünwald/Giemsa cytospins for morphology analysis of Kasumi-1 cells. **e**, Heatmap of gene expression by RNA-seq after shRUNX1-ETO in Kasumi-1 at day2 and day9 (biological triplicates, n = 5810 genes, cut-off FDR < 0.001,). Gene expression of shRUNX1-ETO Day9 compared to shControl Day2-Day9 and shRUNX1-ETO Day2. **f**, Gene expression of shRUNX1-ETO Day9 compared to shControl Day2-Day9. **g-h**, URE-adjusted peak quantification values by ATAC-seq for PU.1 antisense (AsPr, red) and proximal promoter (PrPr, blue) exhibited as AsPr/PrPr ratio (mean value ± s.e.m.) for **g**. RUNX1-ETO knockdown in Kasumi-1 cells (2 days after lentiviral transduction, n = 3) and for **h**. AI-10-49 inhibitor treatment of ME-1 cells (6 hours after treatment, n = 2). **i**, PU.1 asRNA and mRNA transcript expression after shRUNX1-ETO in Kasumi-1 cells (mean value ± s.e.m., n = 6, 2 days after lentiviral transduction) and AI-10-49 inhibitor treatment of ME-1 cells (mean value ± s.e.m., n = 3, 6 hour treatment). **j**, Ratio of PU.1 asRNA and mRNA transcript expression after shRUNX1-ETO in Kasumi-1 cells and AI-10-49 inhibitor treatment in ME-1 cells. **k**, Aligned reads of PRO-seq with quantified peaks for PU.1 AsPr, PrPr, and AsPr/PrPr ratio (right, mean value ± s.e.m.) after shRUNX1-ETO in Kasumi-1 cells (2 days after lentiviral transduction, n = 2). **l**, Aligned PRO-seq reads after RUNX1-ETO knockdown start at the RUNX binding site in PU.1 AsPr.

The −17kb URE creates a regulatory loop with PU.1 PrPr for mRNA expression^2^. Interestingly, the LMPP population presented increased chromatin accessibility at the −17kb URE region (**Extended Fig.4a-b**). We hypothesized that a shift in loop formation towards URE and AsPr instead of PrPr might account for the shift from sense towards antisense transcription in T-cell progenitors (**Fig.1e-f**). To test this idea, we examined chromosomal conformation capture sequencing (Hi-C) data from early T-lymphoid (Jurkat)^19^ and myeloid (HL-60)^20^ cell lines and analyzed PU.1 locus loop formations. While T-lymphoid cells demonstrated an interaction of the URE with intron 3 at the AsPr region, myeloid cells displayed an open locus with an URE-PrPr interaction (**Fig.4a**). We therefore performed in-depth analyses of interacting PU.1 chromosomal structures in both cell lines using chromosomal conformation capture assays (3C) (**Fig.4b**). Quantification of URE to AsPr (URE-AsPr) and URE to PrPr (URE-PrPr) cross-linking frequencies as well as quantification of PU.1 sense and antisense transcripts demonstrated increased URE-AsPr cross-linking frequencies and increased PU.1 asRNA expression in T-lymphoid Jurkat cells (**Fig.4c**). In contrast, in myeloid HL-60 cells URE-PrPr cross-linking frequency and PU.1 mRNA expression were increased (**Fig.4d**). These data suggest a model of two competing promoters for either T-lymphoid or myeloid differentiation (**Fig.4e**). To investigate if RUNX1-ETO plays a direct role in regulating the interplay between the two promoters and the URE, we examined chromatin immunoprecipitation sequencing (ChIP-seq) data^21^. RUNX1-ETO strongly binds to both the AsPr and the URE, with binding decreasing after RUNX1-ETO knock-down (**Fig.4f**) confirming that RUNX1-ETO binds both regions directly through conserved RUNX sites^22,23^ (**Fig.1p**). We further hypothesized that the oncogenic fusion protein could promote a T-lymphoid like chromosomal conformation in which the URE interacts with the AsPr. We thus evaluated if RUNX1-ETO depletion could revert a myeloid state on chromosomal conformation capture sequencing using promoter capture hi-C (CHIC) for enriched annotated promoters^24^. Indeed, a flip towards an increased URE-PrPr interaction was detected (**Extended Fig.4c**). This idea was confirmed by 3C chromosomal conformation analysis showing that after RUNX1-ETO depletion URE-AsPr cross-linking frequencies decreased while URE-PrPr increased (**Fig.4g**) with a concomitant decrease in PU.1 asRNA and an increased in PU.1 mRNA levels (**Fig.4h**). Our data demonstrate that t(8;21) cells adopt a T-lymphoid like interaction state thus switching from PU.1 mRNA towards antisense transcription (**Fig.4i**).

**Fig. 4.**
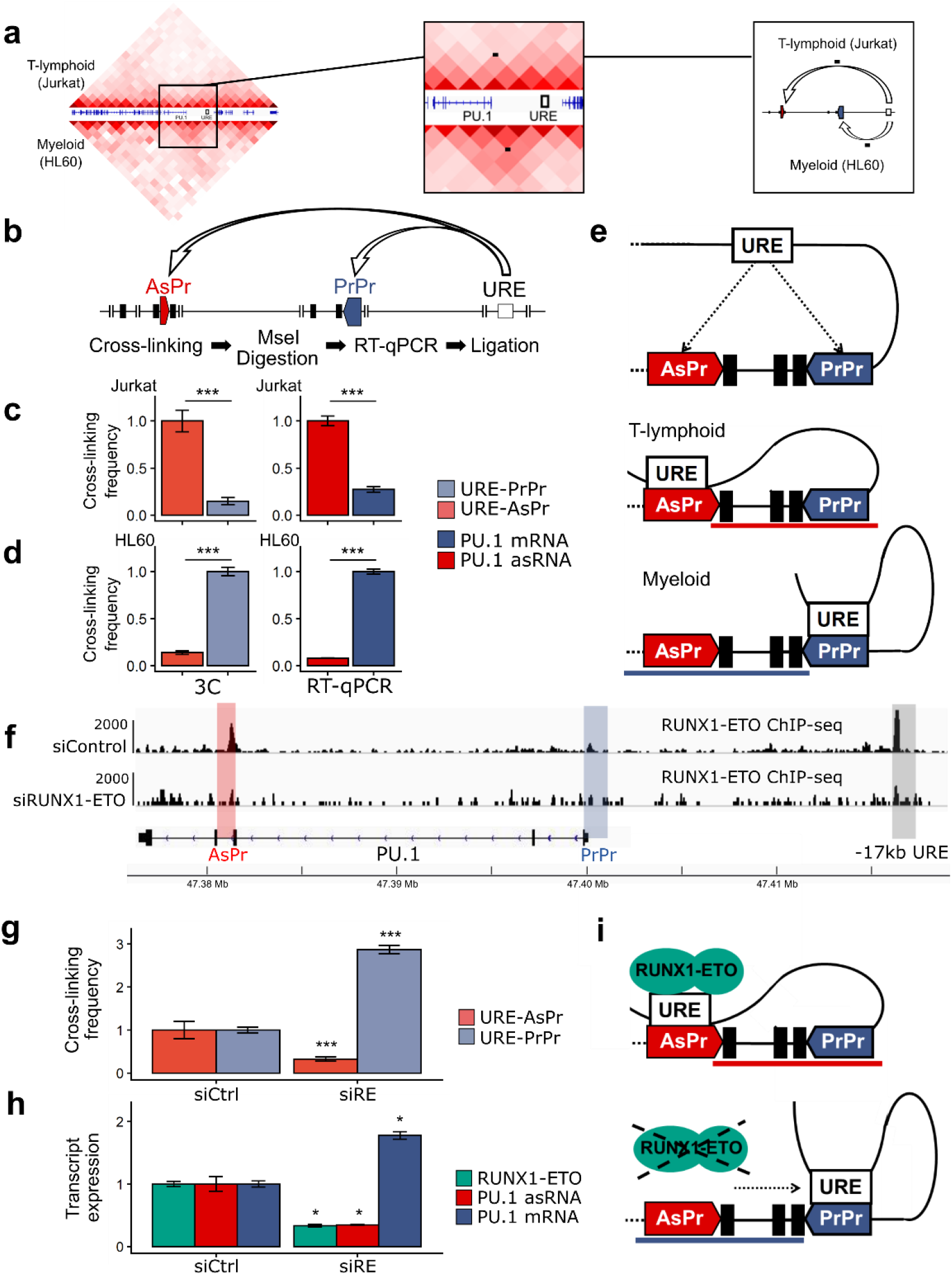
RUNX1-ETO induces a T-like chromosomal conformation. **a**, Chromosomal conformation capture sequencing (Hi-C) in Jurkat (upper half) and HL-60 (lower half) cell lines (B Lucic et al. and Y Li et al.). Black dots indicate the chromosomal looping. **b**, Schematic representation of chromosomal conformation capture assay (3C) for proximity quantification of indicated elements. **c-d**, PU.1 locus chromosomal conformation capture (3C) and transcript expression **c**. in Jurkat and **d**. in HL-60 cells (mean value ± s.e.m., 3C n = 8, RT-qPCR, n = 3). **e**, Model of competing sense/antisense promoters. **f**, Aligned reads of RUNX1-ETO ChIP-seq after small interfering RNA knockdown of RUNX1-ETO (siRUNX-ETO) and mismatch control (siControl) (A Ptasinska et al.). **g**, 3C in Kasumi-1 after siRUNX1-ETO (siRE) and siControl (siCtrl) (mean value ± s.e.m., *p<0.05, n = 3). Mann-Whitney-U test; siRE group versus siCtrl group (**P < 0.01, ***P < 0.001, n = 8). **h**, Transcript quantification in Kasumi-1 after siRUNX1-ETO and siControl (mean value ± s.e.m., *p<0.05, n = 8) of RUNX1-ETO (green), PU.1 asRNA and PU.1 mRNA. **i**, Model RUNX1-ETO inducing a T-lymphoid chromosomal conformation.

Our findings revealed a key mechanism of CBF leukemias which account for 15% and thus the largest group of human acute leukemias^12,15^. We previously reported that the subversion of terminal myeloid differentiation in CBF leukemias is correlated with low levels of PU.1 expression^23^. Conversely, the introduction of PU.1 into CBF RUNX1-ETO leukemic cells induced cellular differentiation and stopped leukemic outgrowth, although the underlying mechanisms have remained unclear^25,26^. We here provide evidence that CBFs regulate the balance between PU.1 sense and antisense transcription that affects PU.1 levels and thus myeloiesis^7^. In addition to PU.1 antisense transcription interfering with PU.1 translation^11^, our data demonstrate reduced mRNA transcription by re-directing a key enhancer away from the proximal promoter towards a promoter of a long non-coding antisense transcript. Recent global analyses of 2,658 whole cancer genomes revealed the presence of non-coding driver mutations ^27^ whose functional role could be crucial, since 60% of the genome are transcribed^28,29^. The promoter of long non-coding transcript PVT1 was recently reported to directly compete with oncogenic MYC promoter for binding a shared set of enhancers, and mutations in the PVT1 promoter were found in breast cancer and malignant lymphoma^30^. The mechanisms how and if these mutations shift the competition towards the oncogenic MYC promoter have remained unclear. In cancer, oncogenes can be activated through genetic translocations that directly juxtaposition the enhancer to an oncogene such as GFI1 or GFIB in childhood medulloblastoma^31^ or EVI1 in rare cases of leukemia^32^. Our findings extend this promoter competition model to a non-coding promoter and provide an explanation how enhancer-promoter interactions are properly coordinated in time, space and in response to the perturbation of differentiation by oncogenes.

## MATERIAL AND METHODS

### Normal donor samples, AML patient samples and cell lines

Normal donor human bone marrow aspirations, peripheral blood cells and AML patient samples for RT-qPCR assays were obtained from the Division of Hematology at Medical University of Graz, Austria, and the Division of Hematology and Hemostaseology at Medical University of Vienna, Austria. Each sample was freshly sorted after Ficoll-Paque mononuclear cell separation revealing >80% blast cells. Material from patients and healthy donors was obtained with approval from the ethics committees at Medical University of Graz (vote number 26369 ex13/14) and Medical University of Vienna (vote number 1184/2014).

Kasumi-1 cells were grown in RPMI1640 supplemented with 20% fetal calf serum (FCS). HL-60 and Jurkat cells were grown in RPMI1640 supplemented with 10% fetal calf serum (FCS). HEK293T and HEK-LentiX were grown in DMEM supplemented with 10% fetal calf serum (FCS). Kasumi-1, HL-60, Jurkat and HEK293T cell lines were obtained from the DSMZ cell line repository. HEK-LentiX cell line for lentivirus production was purchased from Takara Bio. ME-1 cells were purchased from DSMZ and cultured in RPMI1640 with 20% fetal bovine serum, 25 mM HEPES and 100 U/ml Penicillin/ Streptomycin.

### Flow cytometry

Flow cytometry single-cell suspensions were analyzed by flow cytometry using the following monoclonal antibodies (mAbs) conjugated with fluorescein isothiocyanate (FITC), phycoerythrin (PE), PE-Cyanine7 (PE-Cy7), Peridinin-chlorophyll proteins cyanine5.5 (PerCP-Cy5.5), allophycocyanin (APC), APC-Cy7 and Brilliant Violet 570™ (BV605) obtained from BD Pharmingen (BD) or BioLegend (San Diego, CA): CD3 (FITC, HIT3a), CD3 (PE-Cy7, SK7), CD4 (PE, Leu3a), CD8 (APC-Cy7, RPA-T8), CD10 (PE-Cy7, HI10a), CD14 (FITC, M5E2), CD19 (FITC, SJ25C1), CD19 (APC, HIB19), CD34 (PE, 8G12), CD38 (APC-Cy7, HB-7), CD45RA (BV605, HI100), CD66b (FITC, G10F5), CD123 (APC, 6H6), CD235a (FITC, HI264). Viable cells were identified by 4’,6-diamidino-2-phenylindole (DAPI) exclusion. Cells were analyzed by FACSAria Fortessa flow cytometer (BD Biosciences) or sorted by FACSAria Fusion (BD Biosciences) and MoFlo Astrios EQ (Beckman Coulter). FlowJo (Tree Star) was used for data analysis.

### Reverse transcription quantitative PCR

For reverse transcription quantitative PCR (RT-qPCR), total RNA was extracted using Trizol according to manufacturer’s protocol (Gibco). Quantification of PU.1 mRNA, PU.1 asRNA and GADPH expression in sorted normal bone marrow cells, normal peripheral blood mononuclear cells and AML patient samples was performed using Taqman RT-qPCR RNA-to-Ct 1-Step Kit (Applied Biosystems). GAPDH housekeeping gene was quantified using pre-developped TaqMan human GAPDH endogenous control (Applied Biosystems) according to manufacturers’ protocol. Quantification of RUNX1-ETO for assessing knockdown efficiency was performed using high-capacity RNA-to-cDNA Kit and SYBR qPCR Green PCR Master Mix (Applied Biosystems). Quantification of PU.1 mRNA, PU.1 asRNA and GADPH expression in cell lines was performed using strand-specific reverse transcript SuperScriptIII First-Strand Synthesis System (InVitrogen) and GoTaq Probe qPCR Master Mix (Promega) according to manufacturers’ protocol. GAPDH housekeeping used oligodT as primer for stranded cDNA generation. Primers used are described in Extended Table2.

### Electrophoretic mobility shift assays (EMSA)

Nuclear extracts from Jurkat cells were prepared using a Nuclear Extract Kit (40010, Active Motif), according to manufacturer’s recommendation. EMSA was performed using NUSHIFT kit (2005350; Active Motif), according to manufacturer’s recommendation and as previously reported (Di Ruscio et al.). Supershift analysis was carried out by antibodies to RUNX 1 or RUNX 3 (Active Motif cat# 39300 and 39301, respectively). The gel was exposed to X ray film and/or phosphorimaging screens. Oligonucleotide sequences are described in Extended Table2.

### Luciferase reporter assay

HEK293T cells at 80% confluence in 24-well plate were transfected with pXP2 Firefly luciferase, pRL-CMV Renilla luciferase, CBFb, and increasing RUNX1, RUNX2, RUNX3, RUNX1-ETO or CBFb-MYH11 (+RUNX1) expression plasmids (0, 10, 50, 100 and 200 ng). Luciferase activity was measured 48h after transfection using the Dual-Glo Luciferase Reporter Assay System (Promega) and normalized to Renilla luciferase activity.

### Lentivirus production and gene knockdown

Lentivirus was produced by transfection of HEK293T-LentiX (Takara Bio) using PEI (Polysciences) of shRNA-expressing viral vectors with the packaging plasmids pCMVR8.74 and pMD2.G. Virus-containing supernatants were cleared of cellular debris by 0.45μm filtration, concentrated and mixed with 8 μg/ml polybrene. Target cells were exposed for at least 48h to lentiviral supernatant at a multiplicity of infection (MOI) of 4 before downstream applications. Guide-strand sequences for Renilla (shControl), RUNX1-ETO (shRUNX1-ETO) and PU.1 asRNA (shPU.1as) lentiviral shRNA knockdown (sequences in Extended Table2) were cloned in an optimized ‘miRE’ context containing GFP^33^. For RUNX1-ETO knockdown by small-interfering RNA (siRNA), Kasumi-1 cells were transfected with RUNX1-ETO or mismatch control siRNA (sequences in Extended Table2). Cells were collected 24h after transfection for RUNX1/ETO knockdown efficiency assessment depletion and after an additional 40h for cross-linking or RNA isolation^21^.

### RNA isolation and Northern Analysis

RNA isolation, electrophoresis, transfer and hybridization were carried out as described^34^. Polyadenylated mRNAs were selected according to the MicroPoly(A)Purist^TM^ purification kit (Ambion). Preparation of separate nuclear and cytoplasmic fractions was performed according to the Paris™ kit (Ambion). Northern Quantitative analysis was performed on the Storm Phosphorimager. The antisense-specific probe - mixture of two cloned PCR products are described in Extended Table2.

### RNA-seq

Total RNA was prepared using Trizol (Ambion) and processed for sequencing using the NEBNext rRNA Depletion Kit and NEBNEXT Ultra II Direction RNA Library Prep Kit for Illumina. Raw RNA-seq reads were processed using *Trim Galore!* software (http://www.bioinformatics.babraham.ac.uk/projects/trim_galore/) and aligned to GRCh38 using *STAR*^35^. Differential gene expression between groups was calculated using the R package DESeq2 and a FDR < 0.001 was used as cut-off for differential gene expression. Heatmap data visualization was done using ClustVis^36^. Gene Ontology analyses were performed using Panther^37^. Raw RNA-seq data were deposited in the ArrayExpress database (Accession ID: E-MTAB-9016).

### ATAC-seq

Chromatin accessibility mapping was performed using the ATAC-seq technology. In brief, 10^5^ cells were washed once in 50μl PBS and resuspended in a transposase reaction mix containing 12.5 μl 2 × TD buffer, 2 μl transposase (Illumina), 10.5 μl nuclease-free water and 0.01% NP-40. Tagmentation was performed for 30 min at 37 °C. The optimum number of amplification cycles was estimated by qPCR reaction as previously described^38^. Fragments larger than 1,200 bp were excluded by SPRI size selection. DNA concentration was measured using a Qubit fluorometer (Life Technologies). Libraries were amplified using custom Nextera primers^39^ followed by sequencing using the Illumina HiSeq3000/4000 platform. Raw.fastq files were adapter trimmed using *Trim Galore!* software (http://www.bioinformatics.babraham.ac.uk/projects/trim_galore/) and aligned to GRCh37 using bowtie2^40^. Reads corresponding to mitochondrial DNA were removed using samtools (samtools view -@ 20 -h $i | grep -v MT | samtools sort)^41^. Picard tools (http://broadinstitute.github.io/picard) were used to mark duplicate reads arising during PCR as artefacts of the library preparation procedure followed by duplicate read and multimapper removal by samtools (samtools view -@ 20 -h -b -q 30 -F 1024). Bigwig files were generated using deepTools, peaks were called by MACS2 and quantified using the R package diffbind^42^. Intensity values were adjusted to the PU.1 −17kb URE; a quantification inferior to 1:8^th^ of both AsPr and PrPr was used as a filter for outliers. Raw ATAC-seq data were deposited in the ArrayExpress database (Accession ID: E-MTAB-9021).

### PRO-seq library preparation

Precision nuclear run-on sequencing (PRO-seq) for transcriptionally-engaged polymerases in Kasumi-1 after lentiviral small-hairpin RNA (shRNA) knockdown at a multiplicity of infection (MOI) of 4. Nuclei isolation was performed as described in Y. Zhao and colleagues (Cell Rep, 2016). Briefly, 25 million Kasumi-1 cells were collected 48h after lentiviral transduction for RUNX1-ETO knockdown and resuspended in cold swelling buffer followed by Dounce homogenization. Cells were centrifuged and washed with lysis buffer. Nuclei were centrifuged and washed again with lysis buffer followed by freezing buffer and stored at −80°C. Nuclear run-on (NRO) assays were performed with 2 biotin-11-NTPs (biotin-11-CTP and biotin-11-UTP, Jena Bioscience) as described in D. B. Mahat and colleagues (Nat Protoc, 2016). Briefly, nuclei were thawed on ice while the NRO mix was pre-equilibrated at 30°C. NRO was performed at 30°C, terminated with the addition of Trizol-LS (Ambion) and followed by RNA nuclear extraction. Nuclear RNA was fragmented by base hydrolysis and enriched for biotin-labelled RNA with Streptavidin beads (Dynabeads M280 Streptavidin, InVitrogen). Nascent biotinylated RNA was ligated with 3’-VRA RNA adaptor (5’Phos-GAUCGUCGGACUGUAGAACUCUGAAC-3’invdT). Ligated RNA was purified by a second round of Streptavidin beads enrichment and ligated with 5’-VRA RNA adaptor (5’-CCUUGGCACCCGAGAAUUCCA-3’). Ligated RNA was purified by a third round of Streptavidin beads enrichment followed by a reverse transcription step using SuperScript III First-strand synthesis (Invitrogen). Size and quantity of cDNA library preparation is assessed by PCR and determined the number of cycles for library amplification. After full-scale PCR amplification, library size selection was performed by cutting out the desired size after migration in 8% acrilamid gel electrophoresis. Raw PRO-seq.fastq files were processed and aligned to GRCh37 using the proseq2.0.bsh script from the Danko lab^43^. Processed.bam files were sorted and indexed using samtools followed by strand specific bigwig file generation using deepTools^44^. Raw PRO-seq data were deposited in the ArrayExpress database (Accession ID: E-MTAB-9019).

### Publicly available datasets

Data from the following publically available datasets were processed: GSE74912 (ATAC-seq for hematopoietic stem, progenitor and differentiated cells), GSE108266 (AML patient DNase-seq), GSE29222 (ChIP-seq for RUNX1-ETO), GSE69239 (RNA-seq for thymic progenitor cells), GSE93995 and GSE122958 (Hi-C in HL-60 and Jurkat), GSE117107 (Chi-C after RUNX1-ETO depletion in Kasumi-1). ATAC-seq, DNase-seq and RNA-seq data were analysed as described above, with ATAC-seq and DNase-seq following the same URE normalization. Raw ChIP-seq reads were controlled and adapter trimmed using *Trim Galore!* Software (http://www.bioinformatics.babraham.ac.uk/projects/trim_galore/) and mapped against GRCh37 using bowtie2. Multimappers and reads with bad mapping quality were removed using samtools and peak calling was performed by MACS2. HiC and CHiC data were trimmed using trim option of homerTools followed by alignment to hg19 by bowtie2 and tag directory creation by HOMER. Data were visualized using HOMER’s analyzeHiC and TreeView3 software.

### Statistical analysis

All data are presented as mean± standard error mean (SEM). The sample size was determined holding the probability of a type-I error at α = 0.05. Statistical analyses were performed using GraphPad Prism version 6.0.0 for Windows, GraphPad Software, San Diego, California US (www.graphpad.com) and R, R Foundation for Statistical Computing, Vienna, Austria (www.R-project.org). Student’s t-test and Mann-Whitney U test were used to compare differences between two groups. Kaplan-Meier/log rank for survival probabilities of xenografts. P-value of <0.05 was considered statistically significant.

### Chromosomal conformation capture (3C)

For 3C, 50 million cells were prepared for chromatin cross-linking (HL60, Jurkat, Kasumi-1) and centrifuged for 10 min at 200g. Briefly, cell pellets were then resuspended in 21.7 ml fresh culture medium used for cell growth. 1445μL (microliter) of 16% formaldehyde (vol/vol) was added to cross-link cells, gently mixed by pipetting and incubated at room temperature for 10 min. 1.25ml (milliliter) of 2.5M (molar) glycin was added, incubated for 5 min at room temperature and followed by 15 min on ice to stop cross-linking completely. Cross-linked cells were centrifuged for 10 min at 400g, supernatant was removed and cells were quick-frozen on dry ice.

## ACKNOWLEDGMENTS

The authors acknowledge the Core Facilities of the Medical University of Vienna, Austria. Member of VLSI. The authors acknowledge the Biomedical Sequencing Facility, Center for Molecular Medicine (CeMM), Vienna, Austria. A.D.R was supported by the National Cancer Institute of the National Institutes of Health under Award Number R00CA188595 (the content is solely the responsibility of the authors and does not necessarily represent the official views of the National Institutes of Health); Fondazione Cariplo N. 2016-0476 and the Giovanni Armenise-Harvard Foundation; work in C.Bs lab is supported by a grant from Blood Cancer UK (150001); A.K.E. was supported by the R50 CA211304; P.V. was supported by the Austrian Science Fund, SFB grant F4704-B20. D.G.T. was supported by NIH grants R35CA197697 and P01HL131477; and by the Singapore Ministry of Health’s National Medical Research Council under its Singapore Translational Research (STaR) Investigator Award, and by the National Research Foundation Singapore and the Singapore Ministry of Education under its Research Centres of Excellence initiative.” PBS received funding from Austrian Science Fund (FWF) grants P27132-B20 and the Anniversary Fund of the Oesterreichische Nationalbank (OeNB) grant P15936.

