## supplementary_information for "Core binding factor leukemia hijacks T-cell prone PU.1 antisense promoter"

### EXTENDED FIGURE 1

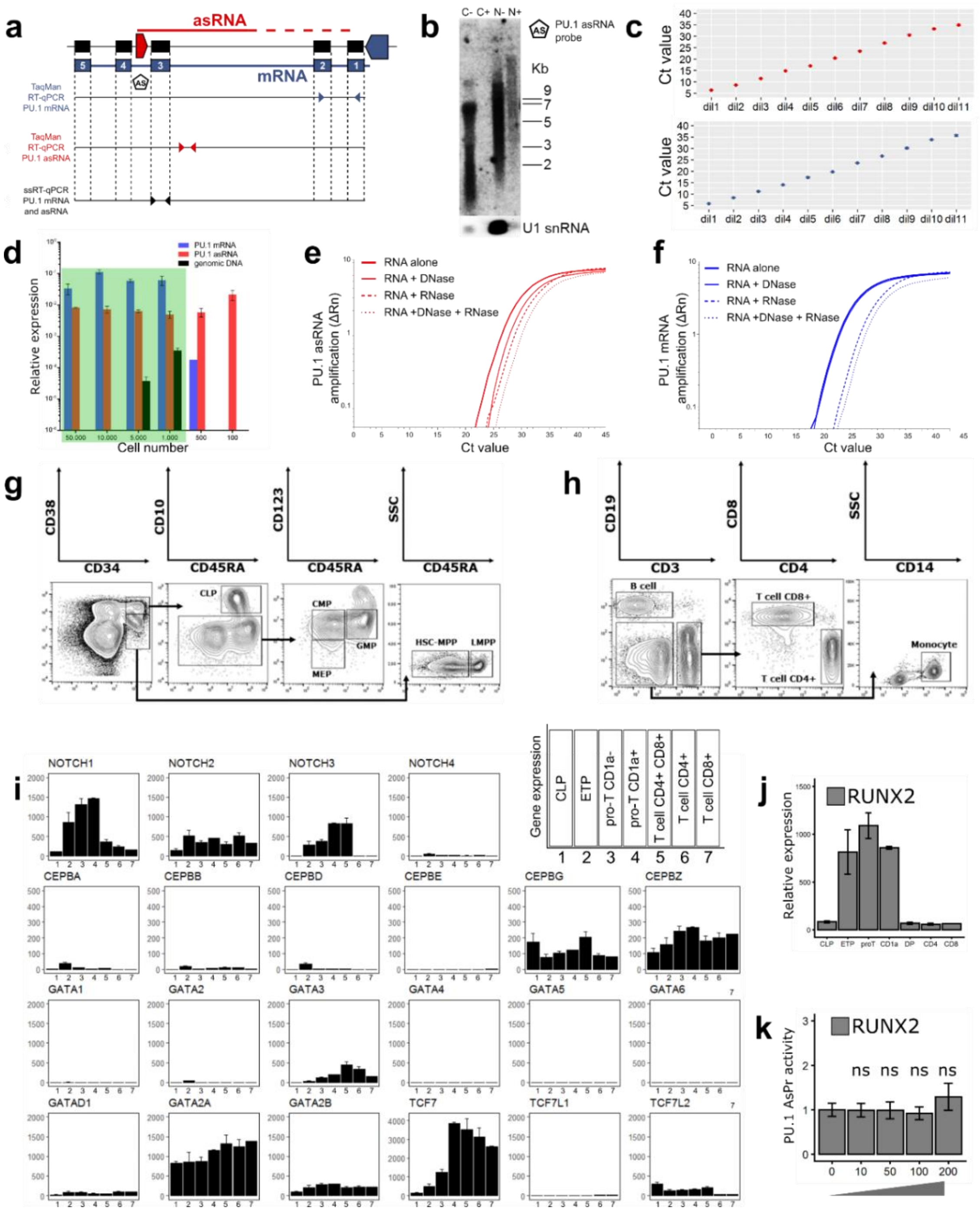


Extended Fig. 1 | **PU.1 quantification assay validation, hematopoietic population isolation for PU.1 transcript quantification and hematopoietic transcription factor mobilization during thymic differentiation. a**, Schematic representation of the PU.1 locus with the proximal (PrPr, blue arrow box) and antisense promoter (AsPr, red arrow box) respectively regulating the transcription of the coding mRNA (blue line, exon number 1-5) and antisense RNA (asRNA, red line). Forward and reverse primer (left and right arrows) pair localization each PU.1 transcript quantification using quantitative reverse transcription polymerase chain reaction (RT-qPCR) or strand-specific RT-qPCR (ssRT-qPCR). Black pentagon arrow shows PU.1 asRNA probe for Northern blot analysis. Colors used in this schematic are consistent throughout the figure. **b**, Northern blot analysis of PU.1 asRNA in HL-60 cell line using PU.1as RNA probe after cytoplasmic (C) or nucleic (N) RNA extraction with (+) or without (-) polyadenylation enrichment. U1 snRNA control probe for nucleic RNA extraction enrichment. Ladder legend for RNA size (Kb, kilobase). **c**, Titration of PU.1 asRNA amplicon (upper panel) and mRNA (lower panel) using the Taqman RT-qPCR assay. Starting DNA at 0.156 ng/µL (dil1, dilution one) is incrementally diluted at a 1:8 ratio (dil2 to dil11, mean value ± s.e.m., n = 2). **d**, Cell limit determination for PU.1 asRNA and mRNA transcript detection using RT-qPCR assay (green area, lowest cell limit, mean value ± s.e.m., n = 2). **e-f**, Characterization of **e**. PU.1 asRNA and **f**. mRNA transcript quantification with RNase and DNase treatments (mean amplification value ΔRn, n = 3) using ssRT-qPCR. **g-h**, Cell isolation using flow cytometry sorting from healthy donors of **g**. total bone marrow (HSC-MPP, merged hematopoietic stem cell and multipotent progenitor; CMP, common myeloid progenitor; MEP, megakaryocyte-erythroid progenitor; GMP, granulocyte-macrophage progenitor; LMPP, lymphoid-primed multipotent progenitor; CLP, common lymphoid progenitor) and **h**. peripheral blood. **i-j,** Gene expression by transcript sequencing (RNA-seq, D Casero et al.) in thymic progenitors and differentiated T cells (ETP, early thymic progenitor) for **i**. most commonly known hematopoietic transcription factors (mean value ± SEM, n=2) and **j**. RUNX2 (mean value ± s.e.m., n = 2). **k**, Luciferase reporter assays in HEK293T cells transfected with PU.1 AsPr reporter plasmid and increasing RUNX2 expression plasmids (plasmid concentration (ng) mean ± s.e.m., n = 4). Student T-test; 0 ng control group versus individual expression plasmid groups (ns, non-significant).

### EXTENDED FIGURE 2

**a**

Detailled cytogenetics and mutations of patient samples from three patient cohorts for PU.1 asRNA and mRNA quantification

| | | cohort1 | cohort2 | cohort3 | |
|---|---|---|---|---|---|
| CBF-AML | t(8;21)(q22;q22) | 5 | 8 | 5 | 36 |
| | inv(16)(p13;q22) | 6 | 7 | 5 | |
| NK-AML | FLT3- NPM1- | 9 | 4 | 5 | 25 |
| | FLT3+ NPM1- | 2 | 1 | - | |
| | FLT3- NPM1+ | 3 | - | - | |
| | FLT3+ NPM1+ | 1 | - | - | |
| BM CD34+ | | 3 | 6 | - | 9 |
| | | | | **Total** | **70** |

**b**

Detailled cytogenetics and mutations of patient samples for DNase-seq PU.1 AsPr and PrPr quantification

| | | | |
|---|---|---|---|
| CBF-AML | t(8;21)(q22;q22) | 3 | 5 |
| | inv(16)(p13;q22) | 2 | |
| NK-AML | FLT3+ | 2 | 14 |
| | FLT3+ NPM1+ | 7 | |
| | NPM1+ | 2 | |
| | CEBPα | 3 | |
| | | **Total** | **19** |

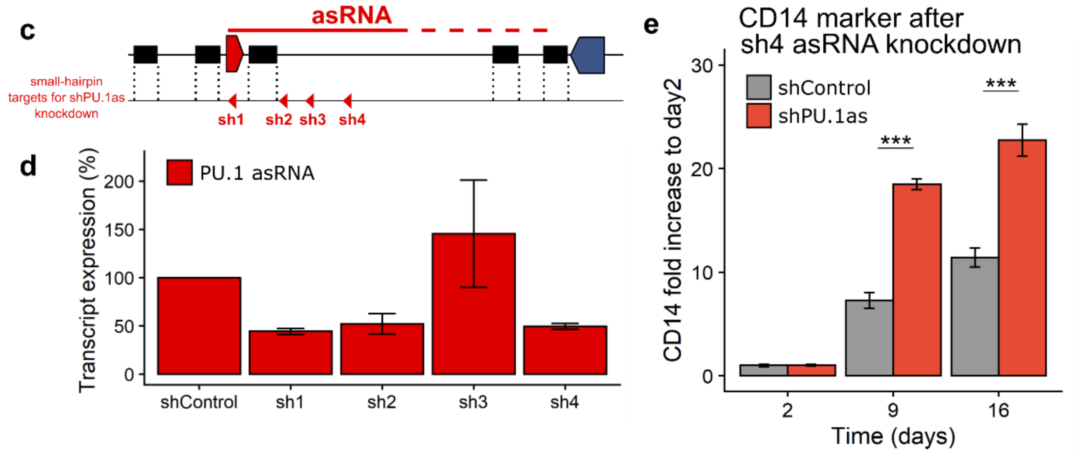


Extended Fig.2 | **Patient data details for core binding factor and normal karyotype AML and lentiviral knockdown of PU.1 asRNA validation. a-b**, Detailed cytogenetic abnormalities and mutations in analyzed patient samples for **a**. PU.1 asRNA and mRNA quantification and transcript ratio calculation and for **b**. PU.1 antisense (AsPr) and proximal (PrPr) promoter DNase hypersensitivity sites (DHS) (SA. Assi et al.). **a**, Core-binding factor AML (CBF-AML) patient group contains the t(8;21)(q22;q22) translocation and inv(16)(p13;q22) inversion anomalies, respectively generating RUNX1-ETO and CBFβ-MYH11 fusion proteins, and is subdivided for each cohort. Normal karyotype AML (NK-AML) patient group is subdivided in Fms-like tyrosine kinase (FLT3) and nucleophospmin (NPM1) mutations, and for each cohort. Bone marrow (BM) CD34+ group contains CD34-enriched bone marrow samples. **b**, CBF-AML patient group contains the t(8;21)(q22;q22) translocation and inv(16)(p13;q22) inversion anomalies and NK-AML patient group contains FLT3, NPM1 or CCAAT/enhancer-binding protein alpha (CEBPα) mutations. **c**, Schematic representation of the PU.1 locus with the proximal (PrPr, blue arrow box) and antisense promoter (AsPr, red arrow box) regulating antisense RNA (asRNA, red line). Small-hairpin targets 1 to 4 (sh1-4) for lentiviral knockdown of PU.1 asRNA (shPU.1as, red arrows). **d**, PU.1 asRNA (red) transcript quantification in Kasumi-1 cells after PU.1 asRNA knockdown (mean value ± s.e.m., n = 2). **e**, CD14 surface marker relative to day2 assessed by flow cytometry after shPU.1as with sh4 target in Kasumi-1 (mean CD14 fold increase relative to day2 ± s.e.m., n = 4).

### EXTENDED FIGURE 3

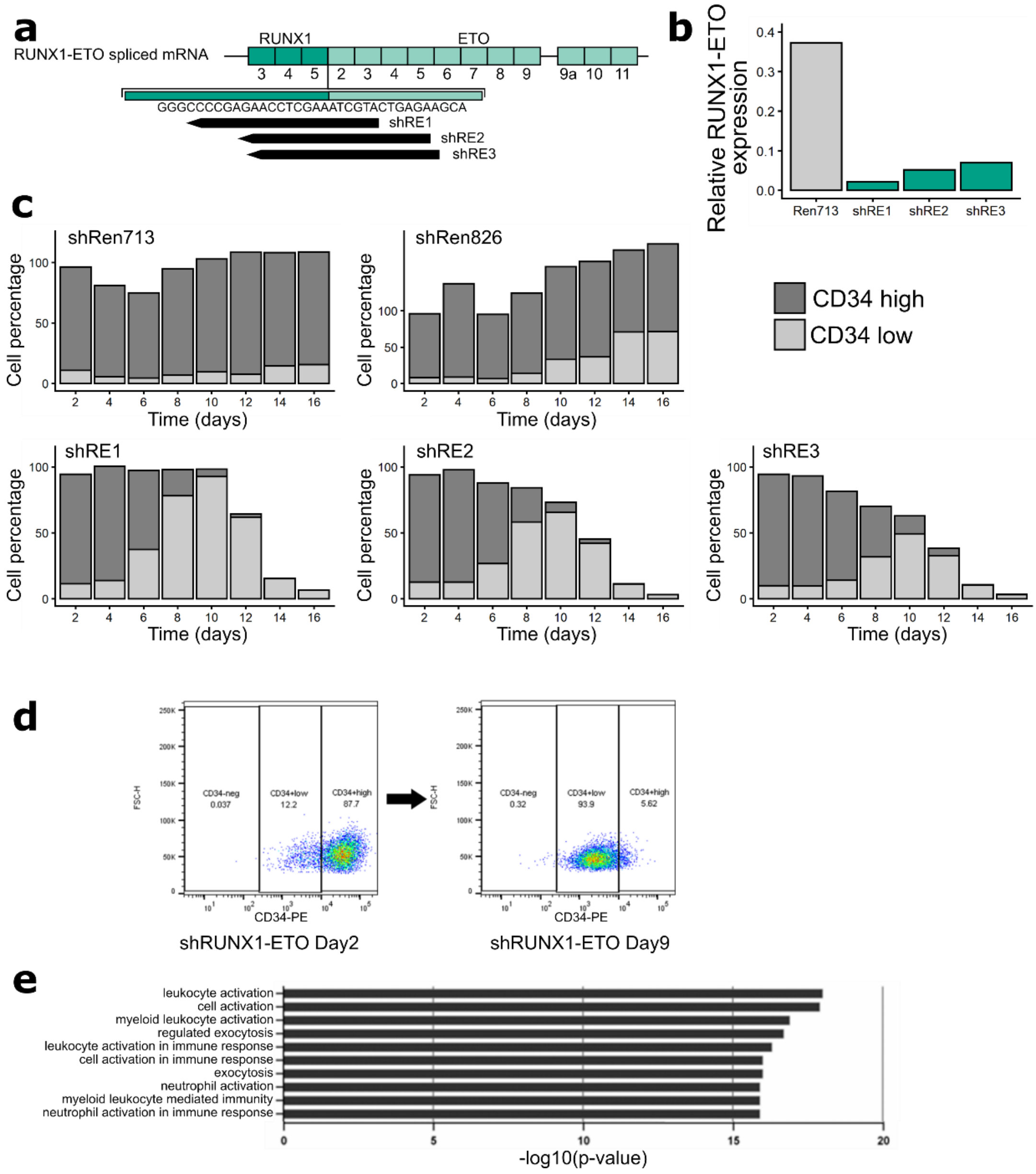


Extended Fig.3 | **RUNX1-ETO depletion by lentiviral small hairpin knockdown prevents cell differentiation and myeloid function. a**, Scheme of shRUNX1-ETO knockdown targets (shRE1-3, small hairpin RUNX1-ETO knockdown for target 1-3). **b**, RUNX1-ETO RT-qPCR relative to GAPDH housekeeping gene after RUNX1-ETO knockdown using each shRE target at day10 in Kasumi-1 cells (n = 1). **c**, CD34 surface marker kinetics assessed by flow cytometry for two negative viability controls (Renilla713 and 826, shRen713 and shRen826) and three constructs for RUNX1-ETO lentiviral knockdown in Kasumi-1 cells (n = 1). **d**, Flow cytometry analysis of CD34 surface marker at day2 and day9 after shRUNX1-ETO (shRE1) in Kasumi-1 cells (n = 2). **e**, Top 10 Pathway gene ontology analysis using Panther from transcript sequencing (RNA-seq) after shRUNX1-ETO in Kasumi-1 cells (values expressed in –log10(p-value), n = 3). Differential expression of shRUNX1-ETO Day9 compared to shControl Day2-Day9.

### EXTENDED FIGURE 4

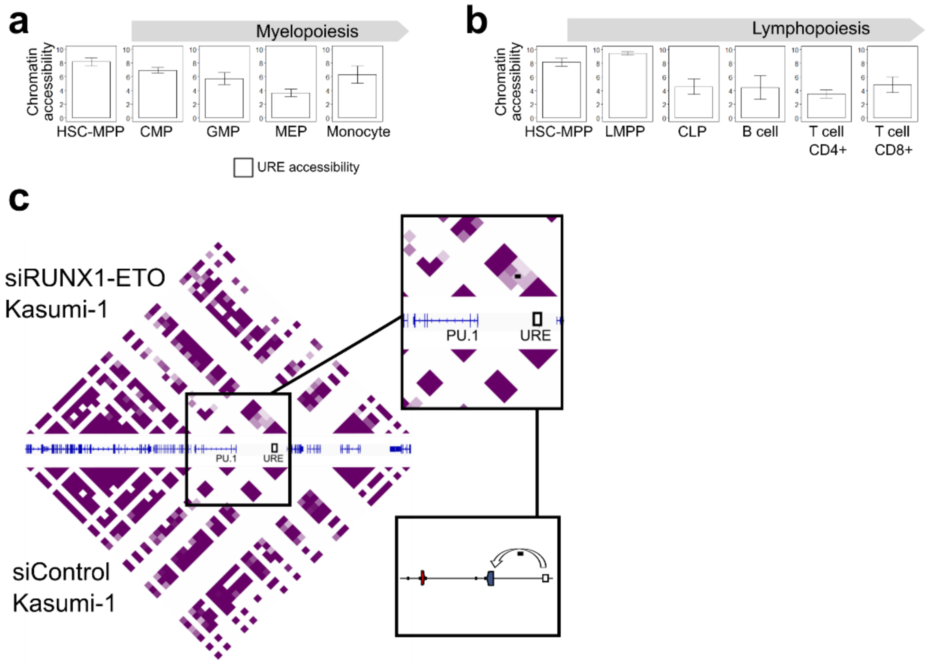


Extended Fig.4 | **PU.1 upstream regulatory element is mobilized in early lymphoid differentiation. a-b**, Quantified area under curve of read alignment peaks for -17kb URE chromatin accessibility using transposase-accessible chromatin sequencing (ATAC-seq) during a. myelopoiesis (HSC-MPP, n=13; CMP, n=8; GMP, n=7; MEP, n=7; Monocyte, n=6) and **b**. lymphopoiesis (LMPP, n=3; CLP, n=5; T cell CD4+, n=5; T cell CD8+, n=5). **c**, Promoter capture chromosomal conformation sequencing (C-HiC) after siRUNX-ETO knockdown. Black dot indicates chromosomal looping of -17kb URE and PrPr (A Ptasinska et al.).

### Bone marrow samples

| Donor: | #1 | #2 | #3 | #4 | #5 | #6 | #7 | #8 | #9 | #10 | #11 | #12 | #13 | #14 | |
|---|---|---|---|---|---|---|---|---|---|---|---|---|---|---|---|
| Age (years): | 65 | 30 | 69 | 66 | 49 | 66 | 69 | 77 | 33 | 68 | 31 | 59 | 58 | 51 | |
| Sex: | M | F | F | M | F | M | M | M | F | M | F | F | F | M | |
| FACS | | | | | | | | | | | | | | | Total |
| HSC-MPP | 1 | 1 | 1 | 1 | 1 | 1 | 1 | 1 | 1 | 1 | 1 | 1 | 1 | 1 | 14 |
| LMPP | | 1 | 1 | | | | 1 | | 1 | 1 | 1 | 1 | | | 7 |
| CLP | 1 | 1 | 1 | | | 1 | | | 1 | 1 | 1 | 1 | 1 | | 9 |
| GMP | 1 | 1 | 1 | | | 1 | 1 | 1 | 1 | 1 | 1 | 1 | 1 | | 11 |
| CMP | 1 | 1 | 1 | 1 | 1 | 1 | 1 | 1 | 1 | 1 | 1 | 1 | 1 | | 13 |
| MEP | 1 | 1 | 1 | 1 | 1 | 1 | | 1 | 1 | 1 | 1 | 1 | 1 | 1 | 13 |
| RT-qPCR | | | | | | | | | | | | | | | |
| HSC-MPP | - | - | - | 1 | 1 | 1 | - | 1 | 1 | 1 | 1 | - | 1 | 1 | 9 |
| LMPP | | - | 1 | | | | - | | - | 1 | 1 | 1 | | | 4 |
| CLP | 1 | 1 | - | | | 1 | | | 1 | 1 | - | - | - | | 5 |
| GMP | 1 | 1 | 1 | | | 1 | - | 1 | 1 | 1 | 1 | - | 1 | | 9 |
| CMP | - | 1 | - | 1 | - | 1 | - | 1 | 1 | 1 | 1 | - | 1 | | 8 |
| MEP | - | 1 | 1 | 1 | 1 | - | | 1 | 1 | 1 | 1 | 1 | 1 | - | 10 |

### Peripheral blood samples

| Donor: | #15 | #16 | #17 | #18 | #19 | #20 | |
|---|---|---|---|---|---|---|---|
| Age (years): | 31 | 31 | 29 | 31 | 28 | 26 | |
| Sex: | F | F | F | M | M | M | |
| FACS | | | | | | | Total |
| Monocyte | 1 | 1 | 1 | 1 | 1 | 1 | 6 |
| B cell | 1 | 1 | 1 | 1 | 1 | 1 | 6 |
| T cell CD4+ | 1 | 1 | 1 | 1 | 1 | 1 | 6 |
| T cell CD8+ | 1 | 1 | 1 | 1 | 1 | 1 | 6 |
| RT-qPCR | | | | | | | |
| Monocyte | 1 | 1 | 1 | 1 | 1 | 1 | 6 |
| B cell | 1 | 1 | 1 | 1 | 1 | 1 | 6 |
| T cell CD4+ | 1 | 1 | 1 | 1 | 1 | 1 | 6 |
| T cell CD8+ | 1 | 1 | 1 | 1 | 1 | 1 | 6 |

### Cell immunophenotypes

| | |
|---|---|
| HSC-MPP | Lin- CD34+ CD38- CD45RA- |
| LMPP | Lin- CD34+ CD38- CD45RA+ |
| CLP | Lin- CD34+ CD38+ CD45RA+ CD10+ |
| MEP | Lin- CD34+ CD38+ CD10- CD123- CD45RA- |
| CMP | Lin- CD34+ CD38+ CD10- CD123+ CD45RA- |
| GMP | Lin- CD34+ CD38+ CD10- CD123+ CD45RA+ |
| Monocyte | CD3- CD19- CD14+ |
| B cell | CD3- CD19+ |
| T cell CD4+ | CD3- CD19- CD4+ |
| T cell CD8+ | CD3- CD19- CD8+ |

Extended Table.1 | Normal bone marrow and peripheral blood donor information, cell sorting strategies, and cell immunophenotypes.

| Sequence name | Sequence (5'-3') | Method |
|---|---|---|
| Target for shRUNX1-ETO | TACGATTTCGAGGTTCTCGGG | shRNA knockdown |
| Target for shPU.1as | GCAGGATCCATTCGGCATTAT | shRNA knockdown |
| Target for shControl | TAGATAAGCATTATAATTCCT | shRNA knockdown |
| Target for siRUNX1-ETO (sense) | CCUCGAAAUCGUACUGAGAAG | siRNA knockdown |
| Target for siRUNX1-ETO (antisense) | UCUCAGUACGAUUUCGAGGUU | siRNA knockdown |
| Target for siControl (sense) | CCUCGAAUUCGUUCUGAGAAG | siRNA knockdown |
| Target for siControl (antisense) | UCUCAGAACGAAUUCGAGGUU | siRNA knockdown |
| FW primer PU.1 mRNA | TGTTACAGGCGTGCAAAATGG | Taqman RT-qPCR |
| RV primer PU.1 mRNA | TGCGTTTGGCGTTGGTATAGA | Taqman RT-qPCR |
| Probe PU.1 mRNA | FAM-AAGACCTGGTGCCCTATGACACGG-TAMRA | Taqman RT-qPCR |
| FW primer PU.1 asRNA | GGTGGCCCTTCCTCTTCAA | Taqman RT-qPCR |
| RV primer PU.1 asRNA | AAGTCTCAGGTCCCAGGAACCT | Taqman RT-qPCR |
| Probe PU.1 asRNA | FAM-TGACCTTAAGCAGTGGCACCTGTGTTCC-TAMRA | Taqman RT-qPCR |
| FW primer RUNX1-ETO | ATGACCTCAGGTTTGTCGGTCG | SYBR Green RT-qPCR |
| RV primer RUNX1-ETO | TGAACTGGTTCTTGGAGCCTCCT | SYBR Green RT-qPCR |
| FW primer GAPDH | GAGTCAACGGATTTGGTCGT | SYBR Green RT-qPCR |
| RV primer GAPDH | AATGAAGGGGTCATTGATGG | SYBR Green RT-qPCR |
| FW primer ssRT-qPCR | AGAGCTTCGCCGAGAACAAC | Taqman ssRT-qPCR |
| RV primer ssRT-qPCR | GCACCATGGGGGTATCGAG | Taqman ssRT-qPCR |
| Probe PU.1 exon3 | FAM-CCGCCACATGGAGCTGGAGC-TAMRA | Taqman ssRT-qPCR |
| RUNX1 complexes oligonucleotide | CCAGGCCTGTGGTGGGGCCCGGGA | EMSA |
| RUNX consensus oligonucleotide | GCAGCTGCATGTCCCAACCACAGCATCC | EMSA |
| Mutated RUNX consensus oligonucleotide | GCAGCTGCATGTCCCAATGGTAGCATCC | EMSA |
| FW intron3 primer | AGGCCTGTGGTGGGGCCCGGGAGC | Northern blot |
| RV intron3 primer | CAGGCGGCAGCCGTTGGAGCTCC | Northern blot |

Extended Table.2 | The sequence list for lentiviral small hairpin knockdown for RUNX1-ETO, PU.1 mRNA and asRNA quantification by RT-qPCR, EMSA and Northern blot analyses.
